# Big trouble with little inserts: boundaries in metagenomic screenings using *lac*Zα based vectors

**DOI:** 10.1101/190520

**Authors:** Luana de Fátima Alves, Tiago Cabral Borelli, Cauã Antunes Westmann, Rafael Silva-Rocha, María-Eugenia Guazzaroni

## Abstract

The vast biochemical repertoire found in microbial communities from a wide-range of environments allows screening and isolation of novel enzymes with improved catalytic features. In this sense, metagenomics approaches have been of high relevance for providing enzymes used in diverse industrial applications. For instance, glycosyl hydrolases, which catalyze the hydrolysis of carbohydrates to sugars, are essential for bioethanol production from renewable resources. In the current study, we have focused on the prospection of protease and glycosyl hydrolase activities from microbial communities inhabiting a soil sample by using the *lacZα*-based plasmid pSEVA232 in the generation of a screenable metagenomic library. For this, we used a functional screen based on skimmed milk agar and a pH indicator dye as previously reported in literature. Although we effectively identified nine positive clones in the screenings, subsequent experiments revealed that this phenotype was not because of the hydrolytic activity encoded in the metagenomic fragments, but rather due to the insertion of small metagenomic DNA fragments *in frame* within the coding region of the *lacZα* alpha gene present in the original vector. We concluded that the current method has a higher tendency for false positive recovery of clones, when used in combination with a *lacZα*-based vector. Finally, we discuss the molecular explanation for positive phenotype recovering and highlight the importance of reporting boundaries in metagenomic screenings methodologies.

## Introduction

Renewable resources, such as plant biomass (essentially lignocellulose), have a significant potential for the production of biofuels and other biotech-produced industrial chemicals due to their higher abundancy and lower price in comparison to other commercial substrates [1]. However, the physicochemical constraints placed on cellulose and hemicellulose polymers by lignin made the saccharification procedure expensive due to a lack of biocatalysts tolerant to process-specific parameters [2,3]. The notorious resilience of bacteria against environmental fluctuations and its inherent biochemical diversity permits screening and isolation of novel enzymes to help overcoming these barriers. Thus, there is a huge amount of gene resources held within the genomes of uncultured microorganisms, and metagenomics is one of the key technologies used to access and explore this potential [4–6].

Functional metagenomics aims to recover genes encoding proteins with a valuable biochemical function [5–7]. For instance, genes considered of interest are enzymes, proteins conferring resistance to diverse physical or chemical stressors, genes coding for catabolic pathways or involved in the production of bioactive compounds, to cite some [47]. The functional metagenomic approach presents two different strategies for libraries generation. Primarily, large-insert libraries, constructed in cosmids or fosmids, allow for the stable recovery of large DNA fragments and sequence homology screening purposes [8]. This strategy would also allow the recovery of complete biosynthetic pathways or the functional expression of large multi-enzyme assemblies (as in the case of polyketide synthases or hydrogenases clusters) [9,10]. On the other hand, small-insert expression libraries (i.e. lambda phage vectors and plasmids), are constructed for activity screening from single genes or small operons [8]. In this strategy, strong vector expression signals (e.g. promoter and ribosome binding site) are used to guarantee that small DNA fragments (2-10 kb) cloned in the vector reach a good chance of being expressed and detected by activity screens [10,11]. At this point, it is of particular relevance mentioning that *lacZα*-based vectors are frequently used in different screenings, with high prevalence in small-insert expression metagenomic libraries [12–17]. In this sense, the blue/white screening inherent of α-based vectors is one of the most common molecular techniques that allows detecting the successful ligation and subsequent expression of the gene of interest in a vector [18–20].

Metagenomics strategies have been of high relevance for providing enzymes used in manufacturing applications [5,7,21]. The use of enzymes in industry has grown considerably, and a number of different categories of enzymes has been used in a wide variety of applications [22]. For example, proteases have been used in detergents, in pharmaceutical and chemical synthesis industries to degrade proteins into amino acids [23]. Glycosyl hydrolases, which catalyse the hydrolysis of carbohydrates to sugars, have been applied to many processes further than bioethanol production (i.e. cellulose and hemicellulose conversion to fermentable sugars), being highly relevant in the textile, paper and food production industries [24].

Studies found in the literature have reported that both enzymatic activities (protease and glycosyl hydrolase) could be found in a single pH based assay using SMA [25,26]. Authors stated that the use of pH indicators dyes such as phenol red or bromophenol blue increases the sensitivity of the assay allowing detection of the acidic shift during hydrolysis of lactose by glycosyl hydrolases (detected as a yellow halo) or casein by proteases (visualized as clear halos) [25,26]. Then, subsequent experiments should be done in order to identify the specific enzymatic activity of the recovered clones [25]. Therefore, in the current study we were interested in obtaining protease and glycosyl hydrolase activities from the microbial community’s inhabitant of a soil sample of a Secondary Atlantic Rain Forest (L. de F. Alves, unpublished results). For this, we have implemented a metagenomic approach using a functional screen based on skimmed milk agar (SMA) and a pH indicator dye (Figure 1A). The metagenomic library was constructed in *Escherichia coli* as a host using the broad host-range vector pSEVA232, which is *lacZα*-based plasmid [27] (Figure 1B).

**Figure 1:**
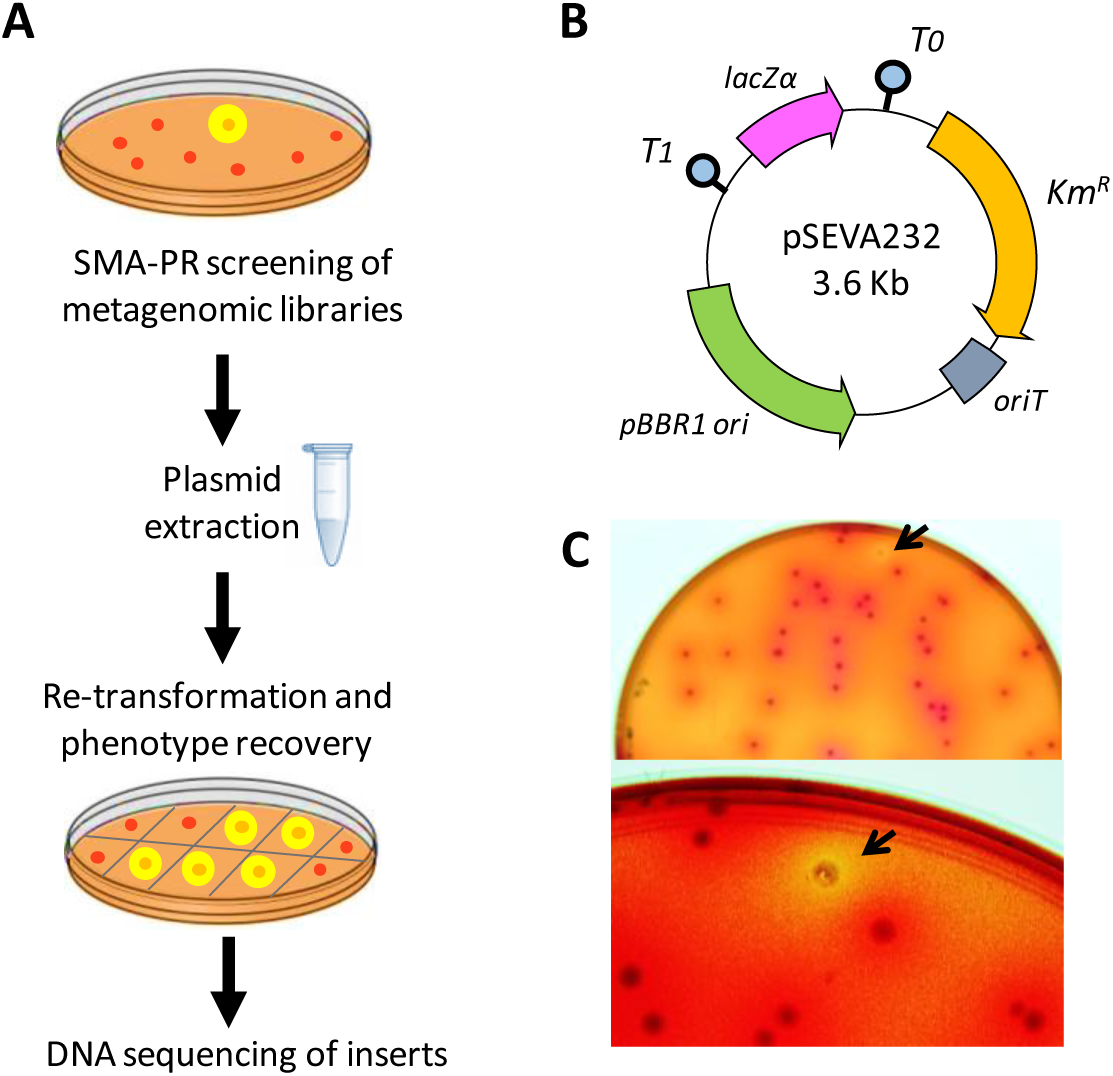
Schematic representation of the workflow for finding novel enzymes (proteases and GHs) using skimmed milk agar (SMA) and phenol red as pH indicator. **(A)** Firstly, metagenomic libraries from soil samples hosted in *E. coli* DH10B were subjected to functional screening in SMA-PR media. Colonies with a yellow halo were taken as potential positive clones, and plasmid was extracted for re-transformation in *E. coli*. Lastly, clones that maintained the phenotype were selected for subsequent restriction profile analyses and sequencing of the metagenomic inserts. **(B)** Overall organization of the structure of pSEVA232 plasmid. Functional elements of the plasmid backbone include an antibiotic resistance marker (*KmR*), a conjugation origin (*oriT*), a broad host-range origin of replication (pBBR1) and the *lacZα* reporter, which contains a multiple cloning site (MCS) in which metagenomic fragments were placed. The T1 and T0 transcriptional terminators are also showed. **(C)** Plate of SMA-PR media after incubation at 37°C for 24 h. Arrow points to a colony of *E. coli* surrounded by a yellow halo and identified as positive clone.

By implementing the SMA-phenol red (SMA-PR) screening approach, we effectively obtained nine clones that were able to generate the typical yellow halos indicative of glycosyl hydrolase (GH) production - although no clear halos, indicative of protease activity, were obtained. However, subsequent experiments revealed that this phenotype was not because of exogenous genes providing hydrolytic activity in these clones. Unexpectedly, restriction profile analyses and sequencing of metagenomic inserts showed that the metagenomic fragments were too small for encoding enzymes able to display activity. Further analyses showed that the metagenomic DNA fragments were inserted *in frame* with the coding region of the *lacZ* gene present in the original vector (α peptide of the β-galactosidase). We concluded that the current SMA-PR method to obtain proteases and GHs have a higher tendency for false positive clones’ recovery, when used in combination with a *lacZα*-based vector. As these vectors are massively used in screenings of small-insert expression libraries, a robust strategy and previous experimental planning should be done to avoid finding and characterizing false positives clones.

## Materials and Methods

### Bacterial strains, plasmids and general growth conditions

*E. coli* DH10B (Invitrogen) cells were used for cloning, metagenomic library construction and experimental procedures. *E. coli* cells were routinely grown at 37°C in Luria-Broth medium [20]. When required, kanamycin (50 μg/mL) was added to the medium to ensure plasmid retention. Transformed bacteria were recovered on LB (Luria–Bertani) liquid medium for 1 hour at 37°C and 180 r.p.m, followed by plating on LB-agar plates at 37°C for at least 18 hours. Plasmids used in the present study were pSEVA232, pSEVA242 [27] and pSEVA242 bearing a 1.5 Kb insert (this study), corresponding the endoglucanase *cel5A* gene from *Bacillus subtilis* 168 [28].

### Nucleic acid techniques

DNA preparation, digestion with restriction enzymes, analysis by agarose gel electrophoresis, isolation of DNA fragments, ligations, and transformations were done by standard procedures [20]. Plasmid DNA was sequenced on both strands using the ABI PRISM Dye Terminator Cycle Sequencing Ready Reaction kit (PerkinElmer) and an ABI PRISM 377 sequencer (Perkin-Elmer) according to the manufacturer’s instructions.

### Screening of GH and protease activities

The metagenomic library used in this study (named LFA-USP3) was generated previously (L. de F. Alves, unpublished results) from a Secondary Atlantic Forest soil sample collected at the University of Sao Paulo, Ribeirão Preto, Brazil (21°09′58.4″S, 47°51′20.1″W). The library was constructed from a microbial community of a soil bearing specific tree litter composition (*Phytolacca dioica*). Metagenomic DNA was cloned into the pSEVA232 vector, a plasmid able to replicate in different gram-negative bacteria, due to its origin of replication [27]. The metagenomic library LFA-USP3 presented about 257 Mb of environmental DNA distributed into approximately 63.000 clones harbouring insert fragments size ranging from 1.5 Kb to 7.5 Kb, with an average size of 4.1 Kb. Briefly, the total microbial DNA was isolated using the UltraClean Soil DNA Isolation Kit according to the manufacturer’s recommendations. The metagenomic DNA was partially digested using Sau3AI and fragments ranging from 2 to 7 kb were directly selected from an agarose gel and purified for cloning into pSEVA232 vector previously digested with BamHI and dephosphorylated. The resulting plasmids were transformed into *E. coli* DH10B cells by electroporation and the cultures were grown in LB-agar plates containing kanamycin (50 μg/mL) for 18 h at 37°C, in order to amplify the library’s number of clones. Clones from the library were pooled together in LB media containing 20% (w/v) glycerol for storage at ‐80°C.

Screening of GH and protease activities was performed according to Jones and collaborators (2007). The library clones were grown in LB-agar plates containing 1% (w/v) skimmed milk, 0.25 mg/mL phenol red and kanamycin (50 μg/mL) for 24h at 37°C. Positive clones were identified due to the formation of a yellow halo around the colonies. Colonies surrounded by a yellow halo against a red background were identified as GH-positive clones and their plasmids were recovered and verified according their restriction patterns when digested using *Nde*l e *Hind*III. The restriction patterns were analysed in agarose gel 0.8% (w/v).

### *In silico* analysis of DNA inserts and identified protein sequences

Putative ORFs from the small fragment sequences were identified using ORF Finder program, available online in (http://www.ncbi.nlm.nih.gov/gorf/gorf.html). Comparisons between the insert amino acid sequences were performed against NCBI database using BLAST (https://blast.ncbi.nlm.nih.gov/Blast.cgi) alignment. There-dimensional models of the chimeric LacZaα/metagenomic peptides (NS1-NS9) and *α*-peptide LacZ were obtained from the QUARK algorithm server (https://zhanglab.ccmb.med.umich.edu/QUARK/) and images were created with PyMOL (http://www.pymol.org/). Thermodynamic analysis of mRNA secondary structure from the different small DNA inserts were performed using the NUPACK algorithms (http://www.nupack.org/). The free energy of a given sequence in a given secondary structure was calculated using nearest-neighbor empirical parameters [29–31]. For each construct, folding energy of mRNA molecule was calculated from positions ‐4 to +70 nt relative to translation star of *lacZ* gene, considering previous data [32] and positions of the DNA inserts (new DNA sequences started at position +53 nt).

## Results and discussion

### Copy number of plasmids alters β-galactosidase expression and halo detection

Previously to the screening for enzymes in the selected SMA-PR media (Figure 1A), we carried out controls for testing the phenotype of clones carrying pSEVA232, the minimal and modular vector used in the construction of the metagenomic library (Figure 1B). For this, we streaked *E. coli* DH10B cultures carrying pSEVA232, pSEVA242, and pSEVA242 bearing a 1.5 Kb insert within the MCS (multiple cloning site) on SMA-PR plates to obtain single colonies. After incubation of the plates for 24 h at 37°C we observed yellow halos around colonies just as in the clones carrying pSEVA242. These results were expected since pSEVA242 is high copy number plasmid (Table 1), carrying the β-galactosidase *α*-fragment in its backbone [27], which guarantees the proper expression of the Lac*Zα* peptide and subsequent protein complementation. As the SMA-PR media contains lactose, its hydrolysis by LacZ produces an acidic shift detected as a yellow halo (Figure 1A).

**Table 1.** Presence or absence of yellow halos indicative of vector-intrinsic β-galactosidase activity in bacterial clones bearing different plasmids.

| Vector plasmid | Yellow halo* | Copy number (copies per cell) | Origin of Replication |
| --- | --- | --- | --- |
| pSEVA242 | Yes | High (100+) | pRO1600/ColE1 |
| pSEVA242- 1.5 Kb insert | No | High (100+) | pRO1600/ColE1 |
| pSEVA232 | No | Medium (30-50) | pBBR1 |

\*Visualization of a yellow halo around colonies after incubation in SMA-PR plates for 24 h at 37 °C.

As explained by Padmanabhan and collaborators (2016) [33], the molecular mechanism for blue/white screening (that is, recovering of functional β-galactosidase LacZ) is based on a genetic engineering of the *lac* operon in the *E. coli* chromosome (coding for the omega peptide with a N-terminal deletion) combined with a subunit complementation achieved with the cloning vector (coding for the *α* peptide). In this way, tetramerization to produce a functional LacZ enzyme is going to occur only if the *α* peptide, which correspond to the intact N-terminal portion of the omega peptide, is added *in trans* [34]. Thus, plasmid pSEVA242 encodes *α* peptide of LacZ protein, which bears an internal MCS, while the chromosome of the host strain (*E. coli* DH10B) encodes the remaining omega subunit to form a functional β-galactosidase enzyme upon complementation. On the other hand, plasmid pSEVA242 bearing a 1.5 Kb insert within the MCS of *lacZα*-gene did not produce a yellow halo, due to the *α*-fragment was disrupted. Finally, pSEVA232, although also being a *lacZ α*-based plasmid, carries a pBBR1 origin of replication, leading to a medium number of copies of plasmids per cell (Table 1), which does not allow enough expression of *lacZ* for proper phenotype production. This feature was essential for using the broad host-range pSEVA232 vector for library construction.

### Screening for proteases and glycosyl hydrolases in SMA-PR may lead to false positives

In order to search for genes coding for proteases and GHs, we screened a metagenomic library hosted in *E. coli* DH10B, which was previously generated in our laboratory (Figure 1A). The screenings were carried out in SMA-PR media, composed by LB-agar supplemented with kanamicine 50 pg/ml, skimmed milk and a pH indicator, the phenol red, that allows to distinguish between GHs (yellow halos) and proteases (clear halos) activities (Figure 1A). From around 70,000 clones screened, we recovered 280 potential positives clones for GHs, of which, just 9 maintained their phenotype when transferred to a new SMA-PR plate (i.e., colonies with yellow halos; Figure 1C). Re-transformed clones were tested to GH activity in SMA-PR plates and plasmids isolated from the colonies surrounded by yellow halos were digested with *Hind*III and *Nde*l enzymes, which revealed 6 recombinant plasmids with unique restriction patterns (Figure 2). Surprisingly, restriction profiles analyses and sequencing of metagenomic inserts showed that the metagenomic fragments were too small (between 42 and 173 bp) for encoding enzymes able to display activity (Figure 2, Table 2).

**Figure 2:**
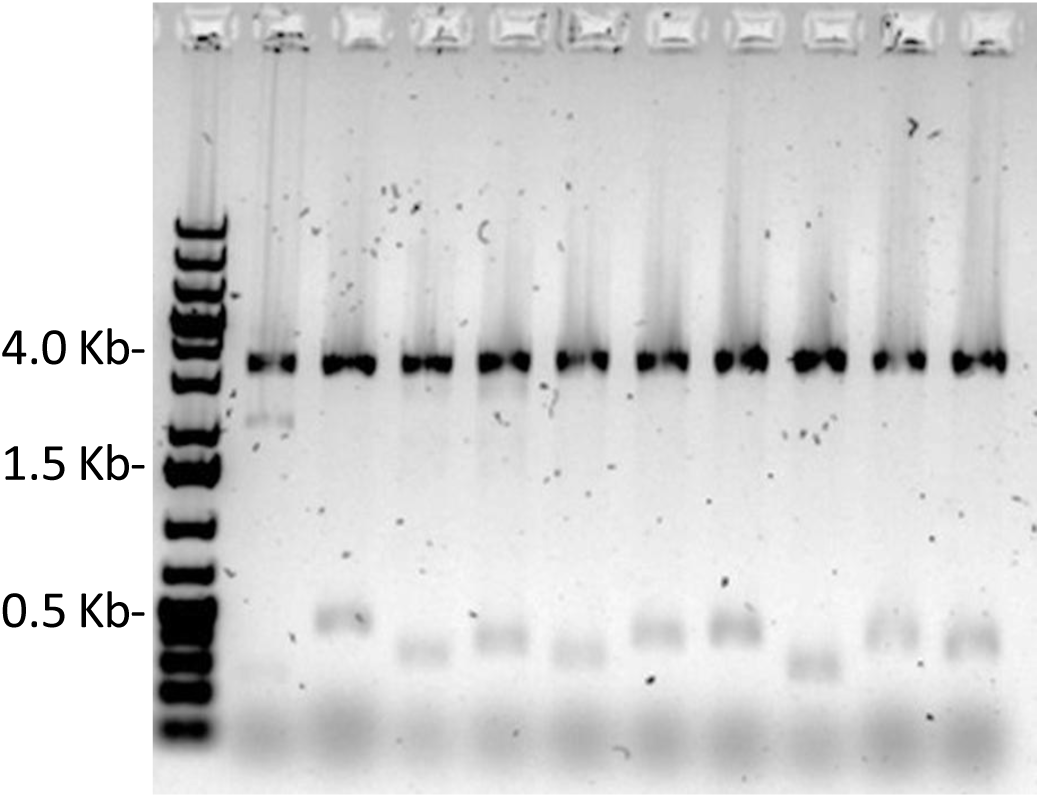
Restriction analysis of plasmids extracted from potential positive clones digested with *Hind*III and *Nde*l in agarose gel 0.8% (w/v). Line 1: empty pSEVA232; lines 2-10: plasmids extracted from clones NS1 to NS9. M: molecular marker GeneRuler 1kb Plus DNA (Thermo Fisher – Waltham, EUA).

**Table 2:**
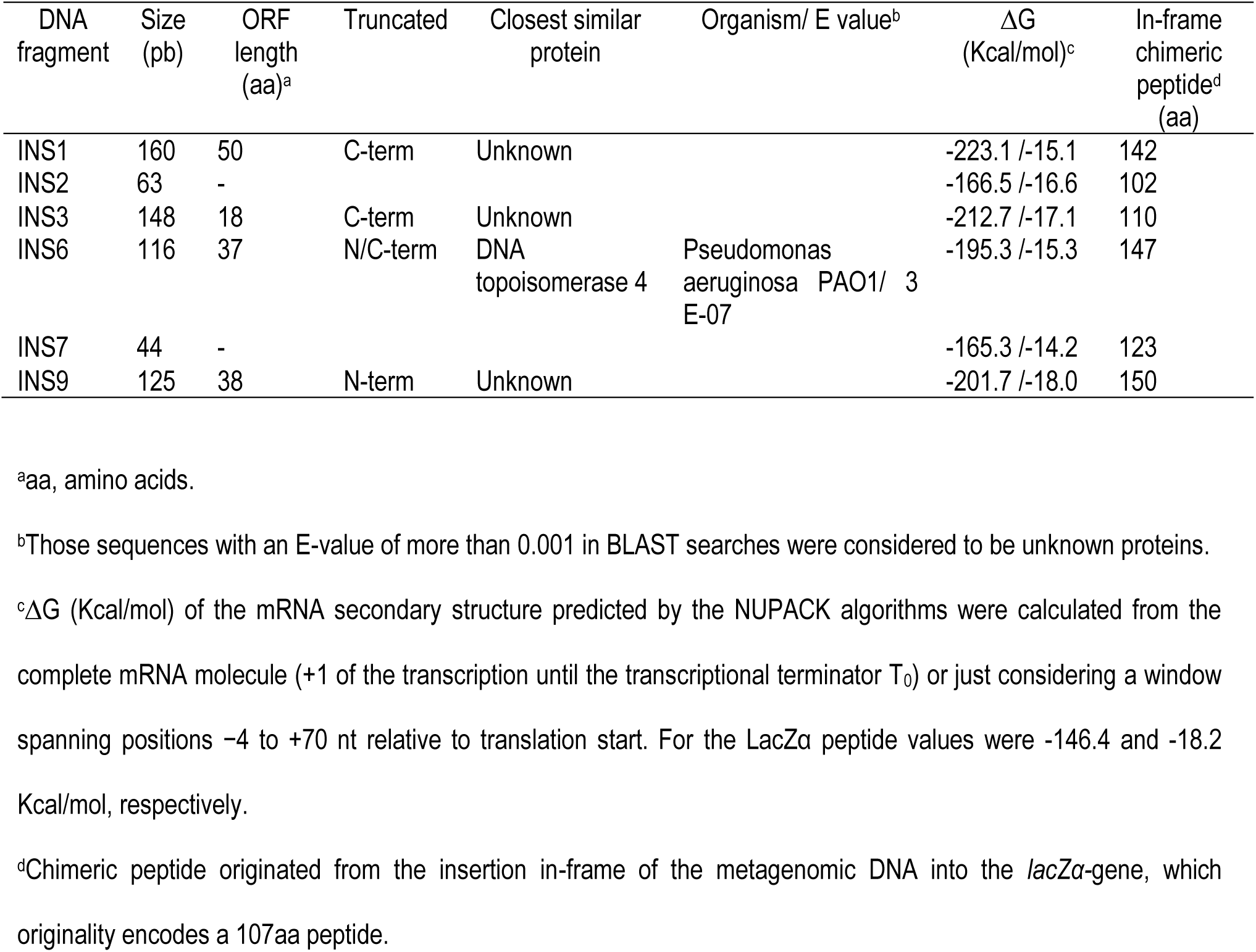
Metagenomic fragments contained in plasmid recovered from positive clones and their sequence features.

*In silico* analysis of the amino acid sequences (Figures 3 and 4) of the chimeric Lac*Zα* fragment/metagenomic peptides resulted from the DNA insertion showed that DNA were inserted *in frame* within the coding region of the *lacZα* gene present in the original vector. Figure 3 shows that complete (DNA inserts NS6, NS7 and NS9) and partial (DNA inserts NS1, NS2 and NS3) recovery of the *LacZα*-peptide were obtained after *in frame* DNA insertion. The N-terminal region of the chimeric *α*-fragment/metagenomic peptides were aligned with the LacZ*α*-peptide looking for conserved amino acids along the N-terminal sequence, although not a clear tendency was observed (Figures 4). On the other hand, three-dimensional modelling analysis of the chimeric peptides in comparison with the original LacZα-peptide resulted in an overall structure maintenance that should assure the activity of the chimeric *α* peptide when is added *in trans* (Figure 5, Figure S1). Taken together, these results indicated that the positives clones were the result of the recovery of functional lacZ*α* polypeptides, showing a strong limitation of the screening technique used.

**Figure 3.**
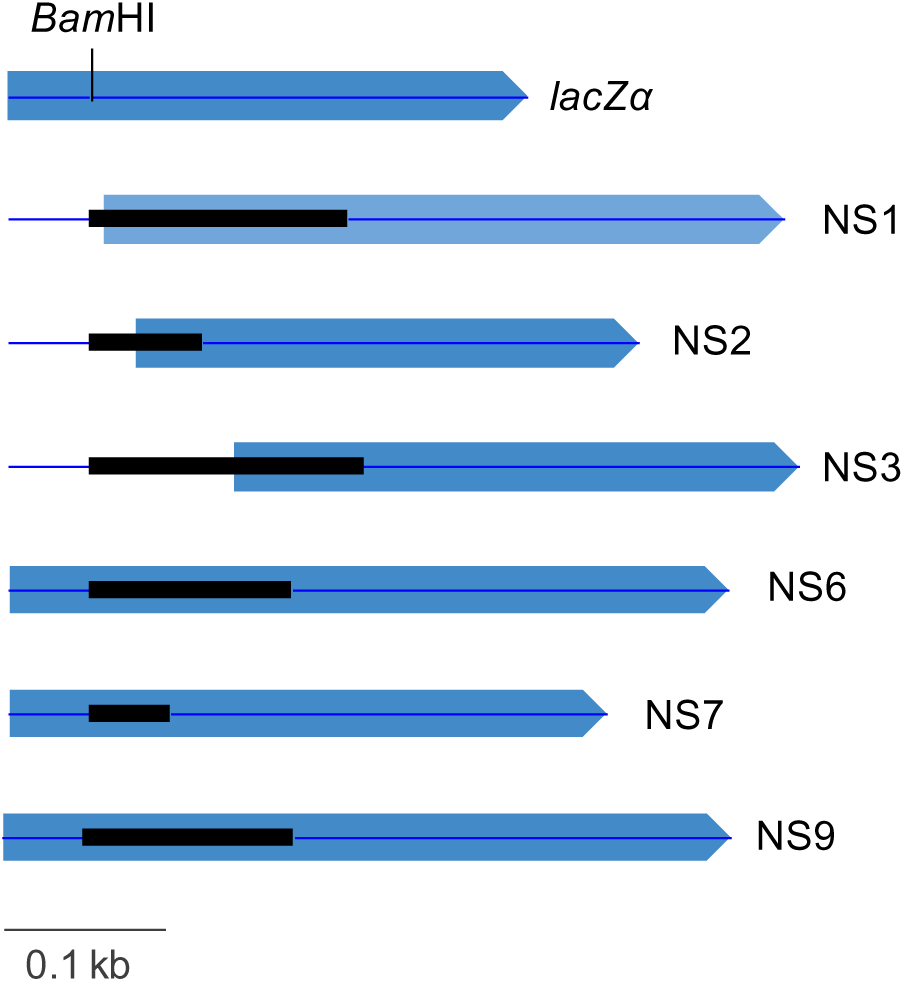
Chimeric LacZα/metagenomic peptides (NS1-NS9) resulted from the *in frame* metagenomic DNA insertion. In blue is shown the DNA sequence coding for the LacZ*α*-peptide and in black the metagenomic insert, cloned in the *Bam*HI restriction site. Complete (NS6, NS7 and NS9) and partial (NS1, NS2 and NS3) recovery of the LacZ*α*-peptide were obtained after *in frame* DNA insertion.

**Figure 4.**
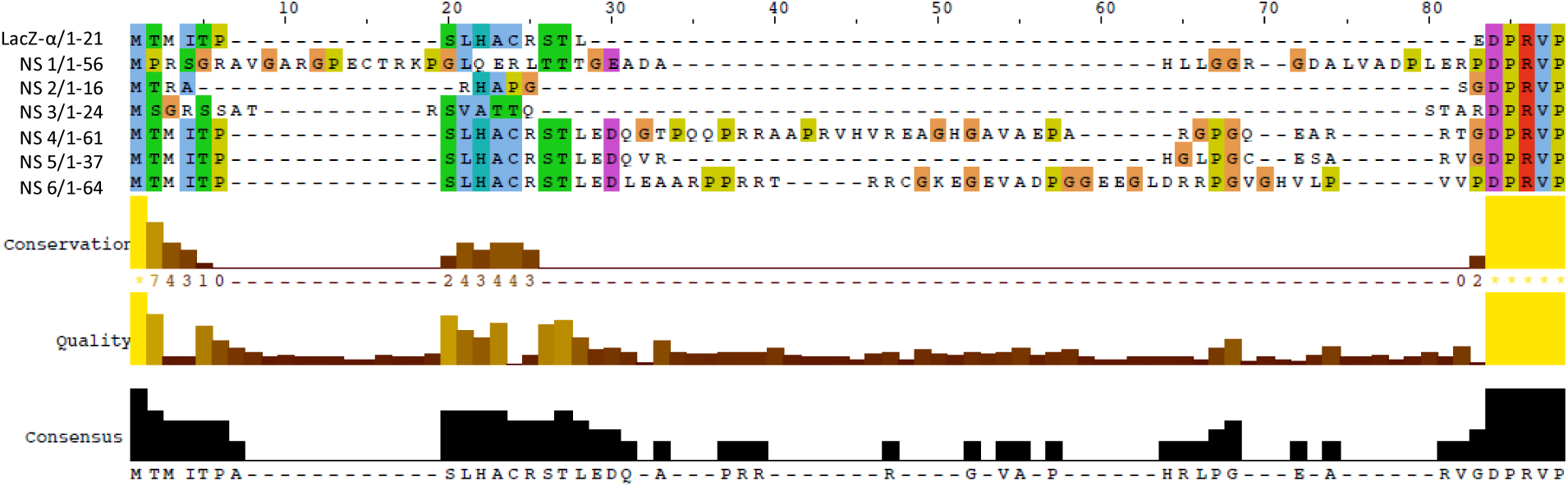
Alignment of the N-terminal region of the chimeric peptides (NS1-NS9) and LacZα-peptide. Alignment was carried out with the T-COFFEE Multiple Sequence Alignment Server [35] and visualization was done with the Jalview program [36]. In general, there were no amino acids conserved along the N-terminal sequence.

**Figure 5.**
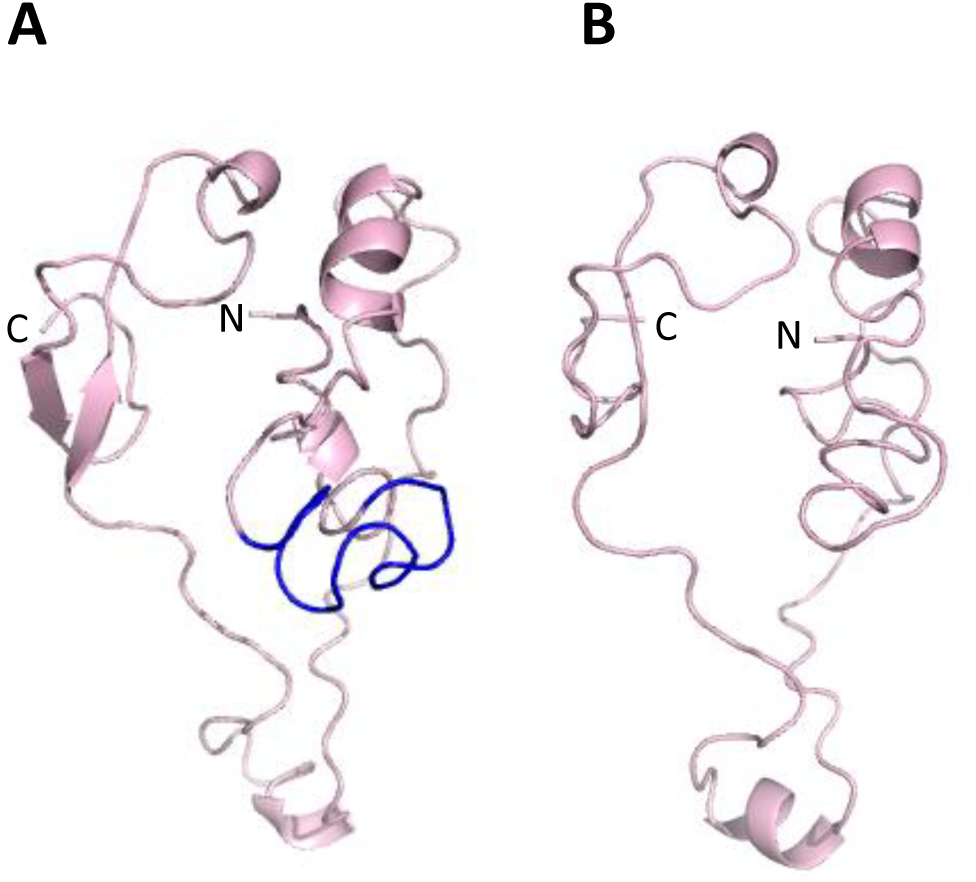
Structural models of the β-galactosidase LacZα peptide (A) and chimeric peptide NS7 (B) resulted from the *in frame* metagenomic DNA insertion. In light pink is showed the 3D structure corresponding to the lacZ*α* peptide and in blue the metagenomic insert. The ITASSER and PyMol softwares were used for structural model’s generation and visualization, respectively [37].

### Reduced free energy in mRNA secondary structure could explain increased expression levels in metagenomic clones

In the light of the evidences presented above, we hypothesized that the recovery of positive clones with very short DNA fragments should be due to the creation of functional lacZα fragments either more active than the original polypeptide or expressed in higher level. In order to elucidate the reason of having identified 9 clones (3 of them repetitive, stressing the intrinsic properties of the clones conducting to that phenotype) from around 70,000 clones screened, that were able to increase the expression of the *lacZα*-gene contained in pSEVA232, we examined in the literature for possible reasons. Preceding studies have shown that the thermodynamic stability of mRNA secondary structure near the start codon can regulate translation efficiency in *E. coli* and other organisms, and that translation is more efficient the less stable the secondary structure [32,38,39]. Although codon bias have been related to slowing ribosomal elongation during initiation and lead to increased translational efficiency [40–42], a recent systematic study using >14,000 synthetic reporters in *E. coli* demonstrated that reduced stability in RNA structure and not codon rarity itself is responsible for expression increases [38]. In this sense, the molecular mechanistic explanation is that tightly folded mRNA obstruct translation initiation, thereby reducing protein synthesis [43].

To get evidence supporting the hypothesis that recovering of the nine positive clones was due to higher expression levels of the chimeric *lacZα*-genes in respect to the original from pSEVA232 (with no phenotype in SMA-PR), we analyzed the local mRNA secondary structure of the different DNA inserts in comparison to the *lacZα* gene. Thus, for each construct (NS1-NS9 and *lac*Z without insert) we computed the predicted minimum free energy ΔG) associated with the secondary structure of its entire mRNA, or the 5′-end region of its mRNA (Table 2). The folding energy of the entire mRNA did not show to be reduced (Table 2). By contrast, the folding energy in position ‐4 to +70 nt relative to translation start showed that in all the new sequences originated by metagenomic DNA insertion the stability of the mRNA molecules was lower than the original, that is, with less negative ΔG values (Figure 6, Table 2). Kudla and collaborators (2009) obtained similar results in respect to the region used for free energy calculation. In this way, studies showed that the region of strongest correlation between folding energy and expression did not overlap with the Shine-Dalgarno sequence [43,44], but with the 30-nt ribosome binding site centered around the start codon [32]. Therefore, results obtained here should explain the identification of the nine clones as positives in the screenings. Consequently, our data are in accordance with previous studies, which demonstrate that reduced mRNAs stability near the translation-initiation site had increased protein expression [32,38,39].

**Figure 6:**
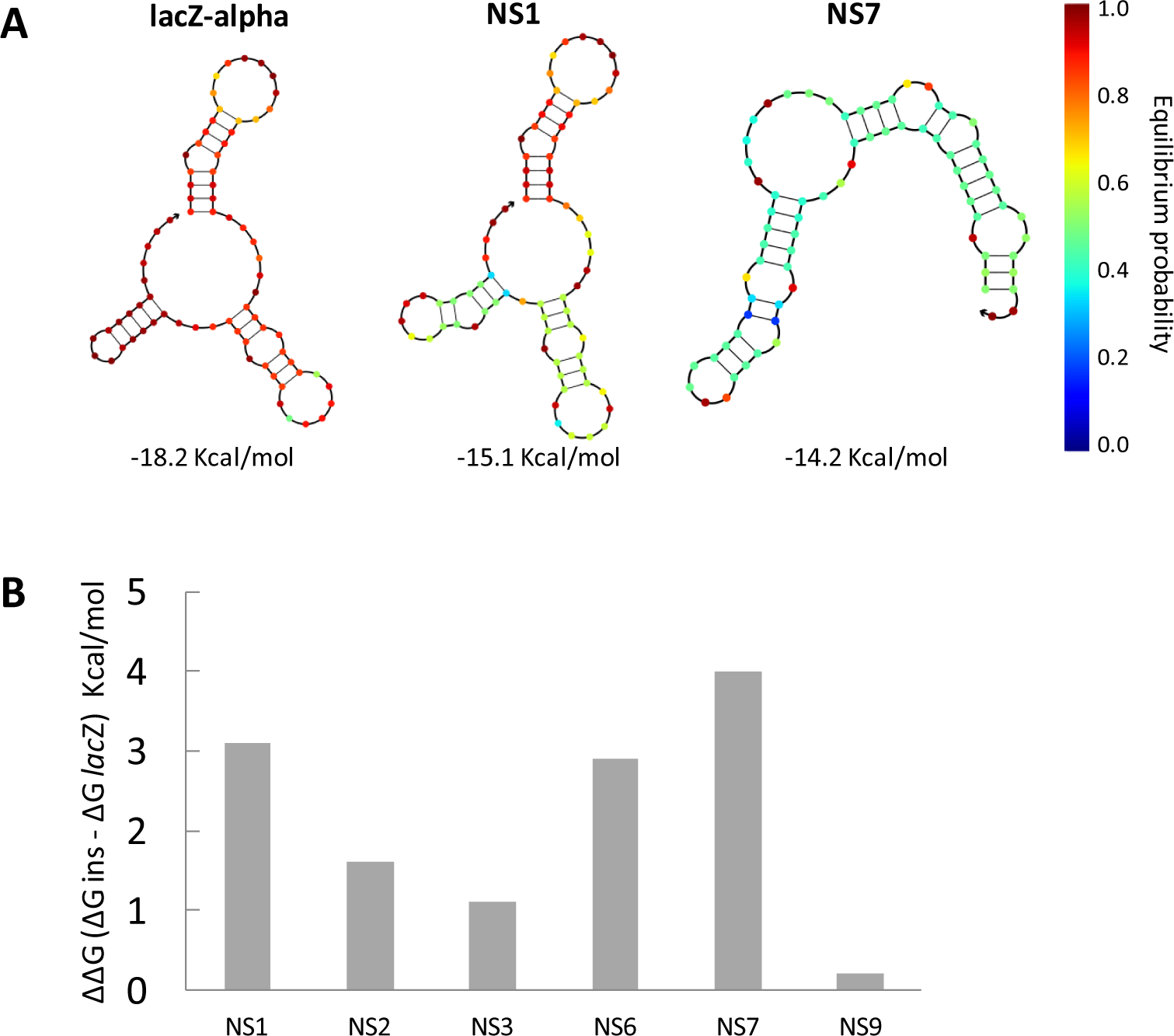
Recovery of clones with halos correlates to relative decreases in free energy of folding when a metagenomic fragment was inserted in the lacZ gene. A) Thermodynamic analysis at 37°C for a dilute solution containing the strand species that interact to form the possible ordered complexes (RNA secondary structures) using the NUPACK algorithms. For each construct, folding energy was calculated from positions ‐4 to +70 nt relative to translation start; three example structures are shown. B) Changes in free energy expressed as ΔΔG considering ΔG values of predicted secondary structure of RNA with (ΔG ins) and without metagenomic inserts (ΔG *lac*Z) expressed in Kcal/mol.

## Conclusions

In the present study, we have used a metagenomic functional approach intending to recover two different types of enzymes in a single assay (i.e., GHs and proteases) using a methodology previously described in the literature [25,26]. For this, we used vector pSEVA232 for library construction, since it displays unique features, such as being minimalist, synthetic, modular and broad host-range [27]. Plasmid pSEVA232 is a *lacZα*-based plasmid, as most of the plasmids used in small-insert metagenomic libraries [12–17]. After the screening in SMA-PR we successfully obtained nine clones showing the typical yellow halos indicative of GH production. However, all were false positive, since small DNA fragments were inserted *in frame* within the *lacZα*-gene present in the original vector. The possible explanation for the positive phenotype of these clones is that the new sequences generated by the metagenomic DNA insertions produced a less stable mRNA molecule at the 5′-end region, which was associated to positively influence protein expression [32,38,39].

Considering that activity-driven screenings (metagenomic or other type, such as in vitro evolution or rational design screens) are time-consuming and use a significant quantity of materials for plate’s preparation, we believe that methods having a higher tendency for false positive clones’ recovery should be avoided. Accordingly, the SMA-PR method when used in combination with *lacZα*-based vectors seems to be extremely inadequate, having high probabilities of obtaining false positive clones. In general, screening strategies using pH based assays, appears to be not sufficiently robust [26]. For instance, we found that microbial secretion, probably derivative from the bacterial metabolism (such as biogenic amines), might act as alkalinizing agents of the solid media, interfering with phenotype visualization [45,46]. We observed that just within a few hours after incubation at 37 °C, halos could turn from yellow to red, hindering clone recovery and reproducibility of the assays. Hence, before embarking on activity-based screenings assays comprising thousands or millions of clones, a robust and straightforward strategy should be planned to avoid finding and characterization of false positive clones. Finally, we strongly encourage the scientific community to report biases in highly accepted protocols, such being the case.

## Acknowledgements

This work was supported by the National Counsel of Technological and Scientific Development (CNPq 472893/2013-0 and 441833/2014-4) and by the Sao Paulo State Foundation (FAPESP, grant number 2015/04309-1 and 2012/21922-8). LFA, TCB and CAW are beneficiaries of FAPESP fellowship (Numbers 2016/06323-4, 2016/06922-5, 2016/05472-6, respectively).

## Supplementary materials

**Supplementary Table 1.**
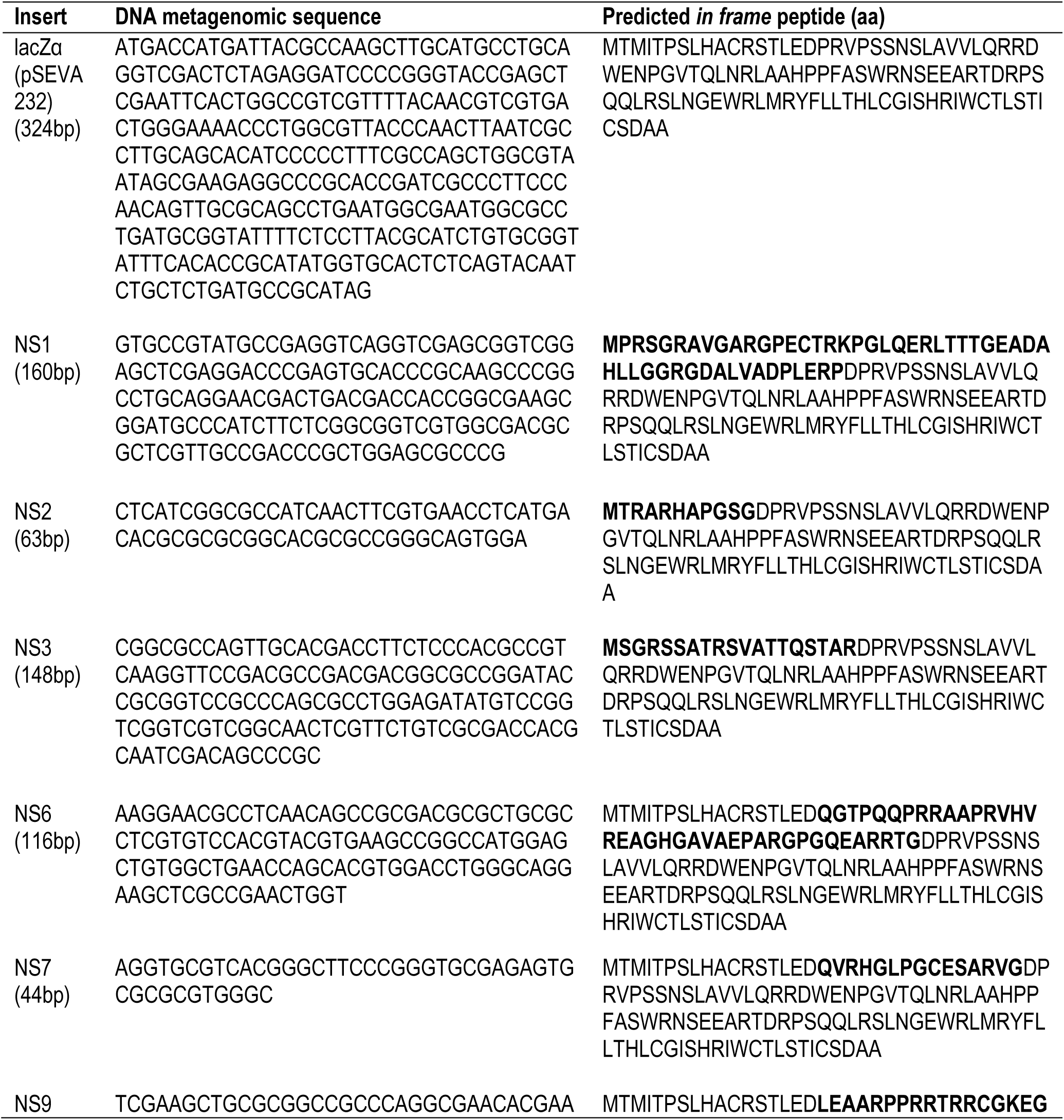
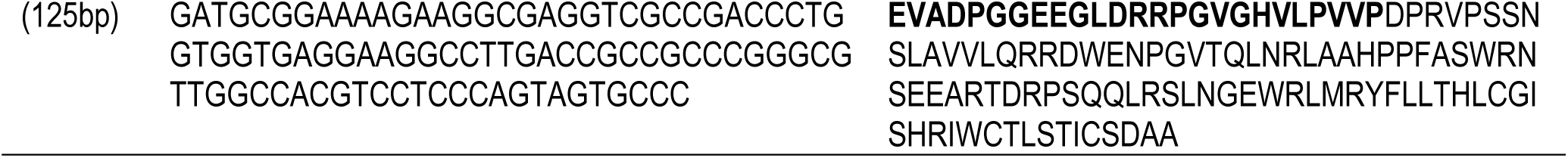
Sequences of the DNA metagenomic inserts derived from the extracted plasmids of potential positive clones. Derivative amino acid sequences of the predicted *in frame* translated peptides are also shown. Amino acids from the metagenomic inserts are in bold.

## References

1. Simmons BA, Loqué D, Ralph J. Advances in modifying lignin for enhanced biofuel production. Curr Opin Plant Biol. 2010;13: 313–320. doi:10.1016/j.pbi.2010.03.001

2. Papoutsakis ET. Reassessing the Progress in the Production of Advanced Biofuels in the Current Competitive Environment and Beyond: What Are the Successes and Where Progress Eludes Us and Why. Ind Eng Chem Res. 2015;54: 10170–10182. doi:10.1021/acs.iecr.5b01695

3. Klein-Marcuschamer D, Oleskowicz-Popiel P, Simmons BA, Blanch HW. The challenge of enzyme cost in the production of lignocellulosic biofuels. Biotechnol Bioeng. 2012;109: 1083–1087. doi:10.1002/bit.24370

4. Dinsdale EA, Edwards RA, Hall D, Angly F, Breitbart M, Brulc JM, et al. Functional metagenomic profiling of nine biomes. Nature. 2008;452: 629–632. doi:10.1038/nature06810

5. Fernández-Arrojo L, Guazzaroni ME, López-Cortés N, Beloqui A, Ferrer M. Metagenomic era for biocatalyst identification. Curr Opin Biotechnol. 2010;21: 725–733. doi:10.1016/j.copbio.2010.09.006

6. Mair P, Gielen F, Hollfelder F. Exploring sequence space in search of functional enzymes using microfluidic droplets. Curr Opin Chem Biol. Elsevier Ltd; 2017;37: 137–144. doi:10.1016/j.cbpa.2017.02.018

7. Lorenz P, Eck J. Metagenomics and industrial applications. Nat Rev Microbiol. 2005;3: 510–6. doi:10.1038/nrmicro1161

8. Danhorn T, Young CR, DeLong EF. Comparison of large-insert, small-insert and pyrosequencing libraries for metagenomic analysis. ISME J. Nature Publishing Group; 2012;6: 2056–2066. doi:10.1038/ismej.2012.35

9. Guazzaroni M-E, Golyshin P, Ferrer M. Analysis of complex microbial communities through metagenomic survey. Metagenomics: theory, methods and applications. Norfolk: Caister Academic Press; 2010. pp. 55–57. Available: https://www.scienceopen.com/document?vid=9f0b077e-dc52-4320-a35f-615c25fc1b55

10. Guazzaroni ME, Silva-Rocha R, Ward RJ. Synthetic biology approaches to improve biocatalyst identification in metagenomic library screening. Microb Biotechnol. 2015;8: 52–64. doi:10.1111/1751-7915.12146

11. Ferrer M, Beloqui A, Timmis KN, Golyshin PN. Metagenomics for mining new genetic resources of microbial communities. J Mol Microbiol Biotechnol. 2008;16: 109–123. doi:10.1159/000142898

12. Mirete S, De Figueras CG, González-Pastor JE. Novel nickel resistance genes from the rhizosphere metagenome of plants adapted to acid mine drainage. Appl Environ Microbiol. 2007;73: 6001–6011. doi:10.1128/AEM.00048-07

13. Lämmle K, Zipper H, Breuer M, Hauer B, Buta C, Brunner H, et al. Identification of novel enzymes with different hydrolytic activities by metagenome expression cloning. J Biotechnol. 2007;127: 575–592. doi:10.1016/j.jbiotec.2006.07.036

14. Guazzaroni ME, Morgante V, Mirete S, González-Pastor JE. Novel acid resistance genes from the metagenome of the Tinto River, an extremely acidic environment. Environ Microbiol. 2013;15: 1088–1102. doi:10.1111/1462-2920.12021

15. Morgante V, Mirete S, de Figueras CG, Postigo Cacho M, González-Pastor JE. Exploring the diversity of arsenic resistance genes from acid mine drainage microorganisms. Environ Microbiol. 2015;17: 1910–1925. doi:10.1111/1462-2920.12505

16. Gao W, Wu K, Chen L, Fan H, Zhao Z, Gao B, et al. A novel esterase from a marine mud metagenomic library for biocatalytic synthesis of short-chain flavor esters. Microb Cell Fact. BioMed Central; 2016;15: 41. doi:10.1186/s12934-016-0435-5

17. Zhou Y, Wang X, Wei W, Xu J, Wang W, Xie Z, et al. A novel efficient β-glucanase from a paddy soil microbial metagenome with versatile activities. Biotechnol Biofuels. BioMed Central; 2016;9: 36. doi:10.1186/s13068-016-0449-6

18. Langley KE, Villarejo MR, Fowler A V, Zamenhof PJ, Zabin I. Molecular basis of beta-galactosidase alpha-complementation. Proc Natl Acad Sci U S A. 1975;72: 1254–7. doi:10.1073/pnas.72.4.1254

19. Zamenhof PJ, Villarejo M. Construction and properties of Escherichia coli strains exhibiting-complementation of ‐galactosidase fragments in vivo. J Bacteriol. 1972;110: 171–178.

20. Ausubel FM, Brent R, Kingston RE, Moore DD, Seidman JG, Smith J a, et al. Current Protocols in Molecular Biology. Molecular Biology. 1994. doi:10.1002/mrd.1080010210

21. Schloss PD, Handelsman J. Biotechnological prospects from metagenomics. Curr Opin Biotechnol. 2003;14: 303–310. doi: 10.1016/S0958-1669(03)00067-3

22. Schoemaker HE. Dispelling the Myths‐‐Biocatalysis in Industrial Synthesis. Science (80-). 2003;299: 1694–1697. doi:10.1126/science.1079237

23. Gupta R, Beg Q, Lorenz P. Bacterial alkaline proteases: Molecular approaches and industrial applications. Appl Microbiol Biotechnol. 2002;59: 15–32. doi:10.1007/s00253-002-0975-y

24. Kirk O, Borchert TV, Fuglsang CC. Industrial enzyme applications. Curr Opin Biotechnol. 2002;13: 345–351. doi:10.1016/S0958-1669(02)00328-2

25. Jones B V, Sun F, Marchesi JR. Using skimmed milk agar to functionally screen a gut metagenomic library for proteases may lead to false positives. 2007;45: 418–420. doi:10.1111/j.1472-765X.2007.02202.x

26. Popovic A, Tchigvintsev A, Tran H, Chernikova TN, Golyshina O V., Yakimov MM, et al. Metagenomics as a Tool for Enzyme Discovery: Hydrolytic Enzymes from Marine-Related Metagenomes. Springer, Cham; 2015. pp. 1–20. doi:10.1007/978-3-319-23603-2_1

27. Silva-Rocha R, Martínez-García E, Calles B, Chavarría M, Arce-Rodríguez A, De Las Heras A, et al. The Standard European Vector Architecture (SEVA): A coherent platform for the analysis and deployment of complex prokaryotic phenotypes. Nucleic Acids Res. 2013;41: 666–675. doi:10.1093/nar/gks1119

28. Santos CR, Paiva JH, Sforça ML, Neves JL, Navarro RZ, Cota J, et al. Dissecting structure-function-stability relationships of a thermostable GH5-CBM3 cellulase from Bacillus subtilis 168. Biochem J. 2012;441: 95–104. doi:10.1042/BJ20110869

29. Serra M, Turner D. Predicting thermodynamic properties of RNA. Methods Enzymol. 1995;259: 242–261.

30. Mathews DH, Sabina J, Zuker M, Turner DH. Expanded sequence dependence of thermodynamic parameters improves prediction of RNA secondary structure. J Mol Biol. 1999;288: 911–940. doi:10.1006/jmbi.1999.2700

31. Zuker M. Mfold web server for nucleic acid folding and hybridization prediction. Nucleic Acids Res. 2003;31: 3406–3415. doi:10.1093/nar/gkg595

32. Kudla G, Murray AW, Tollervey D, Plotkin JB. Coding-sequence determinants of gene expression in Escherichia coli. Science (80-). 2009;324: 255. doi:10.1126/science.1170160.Coding-sequence

33. Padmanabhan S, Banerjee S, Mandi N. Screening of Bacterial Recombinants: Strategies and Preventing False Positives. Mol Cloning - Sel Appl Med Biol. 2011; 3–20. doi:10.5772/22140

34. Gallagher CN, Roth NJ, Huber RE. A rapid method for the purification of large amounts of an alpha-complementing peptide derived from beta-galactosidase (E. coli). Prep Biochem. 1994;24: 297–304. Available: http://www.ncbi.nlm.nih.gov/pubmed/7831210

35. T-coffee: a novel method for fast and accurate multiple sequence alignment1 J Mol Biol. Academic Press; 2000;302: 205–217. doi:10.1006/JMBI.2000.4042

36. Waterhouse AM, Procter JB, Martin DMA, Clamp M, Barton GJ. Jalview Version 2‐‐a multiple sequence alignment editor and analysis workbench. Bioinformatics. 2009;25: 1189–1191. doi:10.1093/bioinformatics/btp033

37. Zhang Y. I-TASSER server for protein 3D structure prediction. BMC Bioinformatics. 2008;9: 40. doi:10.1186/1471-2105-9-40

38. Goodman DB, Church GM, Kosuri S. Causes and Effects of N-Terminal Codon Bias in Bacterial Genes. Science (80-). 2013;342: 475–479. doi:10.1126/science.1241934

39. Gu W, Zhou T, Wilke CO. A universal trend of reduced mRNA stability near the translation-initiation site in prokaryotes and eukaryotes. PLoS Comput Biol. 2010;6: 1–8. doi:10.1371/journal.pcbi.1000664

40. Tuller T, Carmi A, Vestsigian K, Navon S, Dorfan Y, Zaborske J, et al. An evolutionarily conserved mechanism for controlling the efficiency of protein translation. Cell. Elsevier Ltd; 2010;141: 344–354. doi:10.1016/j.cell.2010.03.031

41. Li G-W, Oh E, Weissman JS. The anti-Shine–Dalgarno sequence drives translational pausing and codon choice in bacteria. Nature. Nature Publishing Group; 2012;484: 538–541. doi:10.1038/nature10965

42. Pechmann S, Frydman J. Evolutionary conservation of codon optimality reveals hidden signatures of cotranslational folding. Nat Struct Mol Biol. 2012;20: 237–243. doi:10.1038/nsmb.2466

43. Kozak M. Regulation of translation via mRNA structure in prokaryotes and eukaryotes. Gene. 2005;361: 13–37. doi:10.1016/j.gene.2005.06.037

44. de Smit MH, van Duin J. Secondary structure of the ribosome binding site determines translational efficiency: a quantitative analysis. Proc Natl Acad Sci. 1990;87: 7668–7672. doi:10.1073/pnas.87.19.7668

45. Halász A, Baráth Á, Simon-Sarkadi L, Holzapfel W. Biogenic amines and their production by microorganisms in food. Trends Food Sci Technol. 1994;5: 42–49. doi:10.1016/0924-2244(94)90070-1

46. Silla Santos MH. Biogenic amines: their importance in foods. Int J Food Microbiol. 1996;29: 213–31. Available: http://www.ncbi.nlm.nih.gov/pubmed/8796424

47. Alves LF, Silva-Rocha R and Guazzaroni ME. Enhancing Metagenomic approaches through Synthetic Biology. Functional Metagenomics: Tools and Applications, Berlin: Springer Verlag Berlin. pp. 1–14.

